# Simplifying species-interaction models by grouping parameters: optimal groupings differ between effects and responses

**DOI:** 10.1101/2025.01.18.633749

**Authors:** Christopher R. P. Brown, Daniel B. Stouffer

## Abstract

Most ecological models of species interactions require many parameters, making them expensive to fit to experimental or observational data. To reduce the number of parameters, species are often divided into groups *a priori*, for example on the basis of functional or phylogenetic similarity, and species within these groups are assumed to behave identically. Here, we assess the validity of such grouping by deriving the optimal groupings *a posteriori* based on the fit of the Beverton–Holt model to experimental data from six species of *Drosophila* competing in laboratory conditions. We find evidence that grouping can indeed improve the performance (as judged by AICc) of species-interaction models, but we also observe that there is no single optimal grouping. Further, those groupings that did prove beneficial could not have been easily predicted from available species attributes, calling into question the prevailing method of *a priori* species grouping. Lastly, we find that species did not group equivalently in their competitive effect and in their response to competition, invalidating a key assumption often made implicitly when defining species groupings. Our study suggests the need for a different, more data-informed, approach to grouping in species-interaction models.

## 1 Introduction

Estimating the strength of interactions between species is at the core of many questions of interest in community ecology. Such estimates can be used, for example, to predict the stability of novel assemblages of species (Broekman et al. 2019; Maynard et al. 2020), or to assess the possibility for priority effects in species establishment (Ke and Letten 2018). Of growing interest is also the possibility of placing such estimates in a model to forecast community dynamics (Clark et al. 2020). However, experimentally or observationally determining the strength of each species interaction in a model poses a problem due to the sheer number of parameters that potentially must be determined. Even the simplest models with no environmental dependence require a minimum of one parameter to characterise the interaction of each pair of species (Rocha Filho et al. 2005; Novak et al. 2016). Hence the number of parameters scales quadratically with the number of species; at least 100 parameters are required for the interactions in a community of “just” 10 species. When using a response–surface design, determining these parameters experimentally requires at least one experiment per parameter, and preferably many more (Hart et al. 2018). This is understandably expensive and requires substantial effort. As such, model-based predictions of real communities are typically restricted to a small number of focal species (Ovaskainen et al. 2017; Maynard et al. 2020). However, real communities can contain hundreds of species (Cornell 1999), or even thousands for microbial communities assessed by genetic metabarcoding (Sogin et al. 2006; Warton et al. 2015). To bridge this gap between the size of real communities and our ability to parameterise models to predict their interactions, we need an ecologically-defensible way to reduce the number of parameters that predictive models require.

One simplifying approach is to assume that most pairs of species do not interact or interact weakly, and hence to set many of the parameters to zero (Mutshinda et al. 2009; Weiss-Lehman et al. 2022; Buche et al. 2025). However, weak interactions have been shown to be very important to the stability of ecological communities (McCann et al. 1998; Neutel et al. 2002; Wootton and Stouffer 2016). Setting any individual parameter to zero thus risks losing important information; moreover, we often do not know with confidence which parameters can reasonably be set to zero. A second approach abandons fitting individual species-interaction parameters, instead characterising the community by a handful of community-scale statistical properties (Barbier et al. 2018). While this approach has proven successful at predicting community-scale outcomes, such as diversity and stability, it loses the ability to make inferences or predictions about individual species, which may be important, for instance for management. Here we study a third approach that assumes that species fall into groups, such that all species in the same group interact with species in other groups in an identical fashion, thereby collapsing pairwise interaction parameters for all species in each group into sets of common parameters (Ovaskainen et al. 2017).

In ecological contexts, such groups have typically been defined *a priori* by phylogeny at some taxonomic level (e.g. Gross and Edmunds 2015; Martyn et al. 2021; Buche et al. 2025) or by functional (i.e. trait) similarity (e.g. Uriarte et al. 2004; Martorell and Freckleton 2014; Martyn et al. 2021). The argument for grouping based on phylogeny relies on the principle that closely related species will fill similar ecological niches, and hence will interact with other species in a similar fashion (Darwin 1859; Webb et al. 2002; Godoy et al. 2014). Though this is commonly relied upon, even in grouping individuals into species (Martyn et al. 2021; Buche et al. 2025)—and even more so in microbial contexts where phylogenetic boundaries are fuzzier—empirical tests of this principle have found remarkably little support for it (Cahill et al. 2008; Mayfield and Levine 2010; Godoy et al. 2014). On the other hand, functional groupings (such as growth forms or light guild in plants) have also proven to be poor predictors of species interactions (Bret-Harte et al. 2008; Funk et al. 2017). While attempts have been made to define functional groupings that are more useful in a modelling context (Boulangeat et al. 2012), these rely upon measurements of biotic interactions to define such groups. As such, integrating these into species-interaction models leaves us with a circular argument: a functional group is a group of species that interact with others in a similar way, and we therefore assume that all the species in a functional group interact with others in a similar way. In all, we are left with the question of how best to group species in community models, and whether we can do so in a meaningful way using fewer parameters than required for a fully pairwise parameterisation.

To the best of our knowledge, all studies that use grouping to simplify models of competitive interactions have also made one additional, untested assumption: that the same grouping is equally valid for how species affect others as for how species respond to others (e.g. Ives et al. 2003; Kirwan et al. 2009; Martorell and Freckleton 2014; Gross and Edmunds 2015). There is good reason to question this assumption, as it has been recognised for some time that different traits can govern species’ competitive effects than govern their responses to competition (Goldberg and Landa 1991). For resource competition in particular, competitive effects are determined by how the organism affects local resource availability while competitive responses are determined by how the organism is inhibited by depleted resources (Violle et al. 2009). As such, the best groupings of species may be different for each of the effects and responses.

We propose that species might be classified into groups without regard to any *a priori* definition but instead based purely upon which grouping results in the best performance of the thus-grouped species-interaction model. Such data-informed grouping approaches have previously proved successful in other contexts, such as finding modules in networks (Girvan and Newman 2002; Guimerà and Nunes Amaral 2005; Newman 2006). In this study, we investigate the usefulness and accuracy of such an *a posteriori* classification approach using experimental data from a response–surface competition experiment between six different *Drosophila* species provided by Terry et al. (2021), and examine whether the outcome of such classification may reveal anything about the system’s ecology. We fit a species-competition model, using an underlying Beverton–Holt model for the growth rate, to these data to find the groupings that give the best-performing models (as judged by AICc), and we then compare these groupings to similarity based upon a species trait (body size). While doing so, we group species independently with regard to their effects upon other species and their responses to other species to find whether the data-informed groupings differ between these two cases. We also test the same method on an artificially generated dataset of four species, to validate its ability to find groupings that are known to be present.

## 2 Methods

### 2.1 Data

The data used in this study were drawn from the work of Terry et al. (2021). They consist of fecundity measurements—number of surviving offspring—for six species of *Drosophila*, grown in single-species or two-species communities with varying densities of each species. This experimental design corresponds to a response–surface experimental design commonly used to estimate the strength of species interactions (Inouye 2001; Hart et al. 2018; Cameron et al. 2019; Zhang and Kleunen 2019).

#### 2.1.1 Summary of the experimental methodology

The experiment used six species of *Drosophila* from an Australian rainforest community: *D. birchii*, *D. pallidifrons*, *D. pandora*, *D. pseudoananassae*, *D. simulans*, and *D. sulfurigaster*; for more information on these species, see Jeffs et al. (2021) and Chen and Lewis (2024). The experiment took place in vials with a limited amount of medium for egg-laying and larval development; competition was for space on this medium. A fixed number of adult pairs of one or two species was added to each vial and allowed to mate and lay eggs. The number of adults of each species emerging from these eggs was counted.

Experiments occurred for every single-species and two-species combination of the six species, at a variety of densities. The single-species densities were 6, 18, or 30 adults per vial, and each of these was replicated six times per species. The two-species densities were (6,6), (6,12), (6,18), (6,24), (12,6), (18,6), or (24,6) adults per vial (where the paired values give the densities of each of the two species), and each of these was replicated three times per pair of species. Vials with a single founding pair of founding adults were excluded from this study and the original authors’ due to an Allee effect that the model cannot accommodate, leaving 727 observations in all. Further details of the experimental methodology can be found in Terry et al. (2021).

### 2.2 Model

The data provided the change in population size from one generation to the next. Since these were given by discrete counts, one can model them as the outcome of a negative-binomial process with:

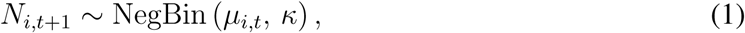

where *N_i,t_*_+1_ is the population at time *t* + 1 (the size of the offspring generation), *µ_i,t_* is the mean of the negative-binomial distribution, and *κ* is an overdispersion parameter.

Following the original authors, we assumed that *κ* was equal for all species. Likewise, we used a version of the Beverton–Holt model to determine the mean given by:

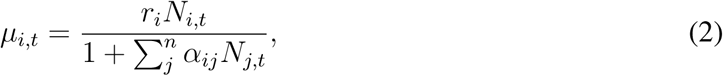

where *N_i,t_*is the population of species *i* at time *t* (the size of the parent generation), *r_i_* is the intrinsic growth rate of species *i*, and *α_ij_* is the per capita effect of species *j* on species *i*. With *n* species, this model has *n* + *n*^2^ + 1 parameters: *n* growth rates, *n*^2^ competition coefficients, and 1 overdispersion parameter. With 6 species, that makes 43 parameters.

#### 2.2.1 Species grouping

In order to reduce the number of free parameters in the model, we grouped species together, and constrained the appropriate competition coefficients to be equal for species within the same group. Within the model defined by eq. 2, each species is effectively in its own group, such that all pairs of species interact in a unique fashion. At the other extreme, all species could be placed in the same group, such that all pairs of species interact equivalently; that is, they are “ecologically equivalent” *sensu* McPeek and Siepielski (2019). Given *n* species, there may be any integer number of groups from 1 to *n*. Groupings are known formally as partitions in mathematics, and the total number of ways to group *n* species is given by the Bell numbers, denoted *B_n_* (Bell 1934).

To allow species to be grouped differently in their competitive responses than their competitive effects, we decoupled the two groupings. Thus, each grouped model was defined by two independent groupings of species: one grouping of the competitive responses and one of the competitive effects. With six species and independently selected row and column groupings, the total number of possible models is (*B*_6_)^2^ = 41 209. The species’ intrinsic growth rates, *r_i_*, were not affected by grouping; each species always had its own growth rate fitted independently of other species. The total number of parameters in a grouped model is thus reduced to 1 overdispersion parameter, *n* growth rates, and a number of competition coefficients equal to that model’s number of row groups multiplied by that model’s number of column groups.

Selecting the groups is equivalent to the machine-learning problem of classification: each species must be placed into one group, or “class”, with other species like it. Our approach is a kind of unsupervised classification, as we have not *a priori* identified the groups, or even how many groups we require (Alloghani et al. 2020). However, it is not fully unsupervised, as we do know to which species each observation belongs, and are thus trying to sort the species, and not the raw observations, into groups.

To better understand the meaning of species grouping of competition coefficients, it is useful to recognise that this component of the model can be represented as an interaction matrix (Novak et al. 2016), where a row contains all the competitive effects on a single species and a column contains all the competitive effects of a single species. If two species are assumed to have the same responses to competition, then two rows of the interaction matrix are constrained to be equal. Similarly, if two species are assumed to have the same competitive effects, then two columns of the interaction matrix are constrained to be equal. This allows us to refer below to the grouping of the competitive responses as the “row grouping” and the grouping of the competitive effects as the “column grouping”. Formally, if species *a* and *b* are in the same row group, *α_aj_* ≡ *α_bj_* ∀*j*, and if *a* and *b* are in the same column group, *α_ia_* ≡ *α_ib_* ∀*i*.

#### 2.2.2 Model fitting

We identified the maximum likelihood parameter values of the models with the SbPlx algorithm of the NLopt C^++^ package (Rowan 1990; Johnson 2007). When fitting the model to the data, the competition coefficients were not constrained to be positive; this allowed for the possibility of facilitative effects between some species, as these have been shown to be important in natural systems (Bimler et al. 2018; Buche et al. 2025). Growth rates and the overdispersion parameter were constrained to be positive.

Just like the intercept and slope in a linear regression, the Beverton–Holt model will exhibit some covariance between estimates of parameters—for instance, all the competition coefficients for the response of a species will be positively correlated with the growth rate of that species. While this will affect the precision of parameter estimates, and thus may affect the accuracy or confidence interval of any predictions made from a parameterised model, it will not affect the likelihood estimates of the models. As we used the likelihood (via AICc, see below), and not the parameter estimates themselves, to determine the optimal groupings, this covariance of parameters should not have impacted the results.

#### 2.2.3 Assessing model performance

To assess the performance of the different species groupings, we exhaustively fit all 41 209 possible models and computed the resulting AICc for each of those fits. As a measure of performance, we selected AIC over BIC on the basis that we want to find the best model for making ecological predictions rather than seeking the “true” model (Aho et al. 2014). Furthermore, AIC is more conservative in promoting grouping as BIC imposes a greater penalty for the number of parameters for any reasonably sized data set. If there is insufficient replication, this may lead to there not being enough data to differentiate two species, even if they are ecologically distinct. Thus, a more conservative approach is less likely to be mistaken. Although the number of observations was large on the whole, some of the models had a relatively high number of parameters compared to the number of observations (727 observations compared to a maximum of 43 parameters), so we used AICc to correct for this “small sample size” effect (Burnham and Anderson 2002). AICc values can be compared to determine which model performs the best, but they cannot directly be interpreted to find the significance of the difference in performance. As such, we also converted the set of AICc values to Akaike weights, which give the probability that each model is the best among the models assessed (Wagenmakers and Farrell 2004).

Using these Akaike weights, we calculated two important statistics. First, the total Akaike weight of models with a lower AICc than the entirely ungrouped model (eq. S13); this is the probability that some model performs better than the model that assumes all species are statistically unique. Second, the total Akaike weight of models that used the exact same grouping for the rows as for the columns (eq. S14); this is the probability that the best-performing model was one that grouped species equivalently in their competitive effects and competitive responses. To evaluate the significance of the latter, we performed a Monte Carlo permutation test, comparing the total Akaike weight of the *B*_6_ = 203 models that grouped rows and columns equivalently to the total Akaike weight of 999 equally sized random subsets of the models. The aim of this test was to determine whether the best model was more likely to be one that grouped rows and columns equivalently than we would expect by chance.

#### 2.2.4 Coclassification matrices

To summarise the results, we also used the Akaike weights to generate two species-by-species coclassification matrices, one for the row groupings and one for the column groupings. Each cell of a coclassification matrix represents the coclassification of a pair of species, and contains the total Akaike weight of models that place those two species in the same group. As the Akaike weight of each model represents the probability that it is the best-performing model, each cell in the coclassification matrix represents the total probability that the best-performing model is one of the models that coclassifies those two species. For reference, note that the diagonal of a coclassification matrix is tautologically equal to 1.

There are several useful reference values for the off-diagonal elements of these matrices. A neutral model approach, where all species are identical, would expect all species to be in the same group: every off-diagonal cell would therefore be 1 (eq. S10). If each species were expected to be unique, every off-diagonal cell would instead be 0 (eq. S11). Alternatively, if every possible grouping were considered to be equally likely, every off-diagonal cell would be 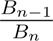 : 0.256 for six species (eq. S12). Unfortunately, without a null distribution of coclassification matrices, it was impossible to test the data-informed coclassification matrices against any of these reference values.

However, we could test the row-grouping and column-grouping coclassification matrices against one another, and against a known species trait: body size (as mean wet mass of female adults, obtained from Terry et al. 2021). Body size is often an important trait in determining the parameters of species-interaction models, as it determines many key physiological rates (Hudson and Reuman 2013) Firstly, we converted the coclassification matrices to dissimilarity matrices by subtracting them from 1. We used PERMANOVA (through the adonis2 function of the vegan R package; Oksanen et al. 2020) to test whether these dissimilarity matrices could be explained by body size (Anderson 2001). This consisted of two PERMANOVA tests, both using the body size for each species as the predictor variable, and using either the dissimilarity matrix for the row groupings or for the column groupings as the response variable. We also performed a Mantel test (Mantel 1967) for correlation between the row-grouping and column-grouping dissimilarity matrices, thus providing a second test of the assumption that it is appropriate to use the same grouping for both. Though the Mantel test can be incorrect when dissimilarity matrices are calculated as distances between points in multivariate space (as it is most commonly used in ecology), with dissimilarity matrices calculated from Akaike weighted coclassification, the dissimilarity matrices themselves are the objects of interest, and the Mantel test remains appropriate (Legendre et al. 2015).

#### 2.2.5 Group resolution

To validate the method’s ability to detect groups when they are present, we employed the same method on simulated data with known groups. We generated simulated data with four species in two equally sized groups with known between group differences in the interaction strengths, and we then used the method to analyse the resulting coclassification from these datasets. The full details of this test are presented in section S1.

As expected, coclassification between species in different groups decreased as the difference in competition coefficients between the groups increased, both for competitive effects and responses (fig. S4). However, this validation also showed that the method has a greater ability to distinguish groups in competitive effects (columns) than in competitive responses (rows) (i.e. coclassification between species known to be in different effect groups decreased more quickly with difference between the groups than did coclassification between species in different response groups). For species known to be within the same group, coclassification probability was not much affected by the difference from other groups, averaging between 0.5 and 0.75 regardless of the magnitude of difference from other groups.

## 3 Results

### 3.1 Model performance

Across all models, the AICc values ranged from 6166.79 to 6278.59. The ungrouped model gave an AICc value of 6176.14. Figure 1 shows the distribution of AICc values from the models, with some models of interest marked. Plots of predicted against observed values for these models of interest are shown in figs. S1–S3.

**Figure 1:**
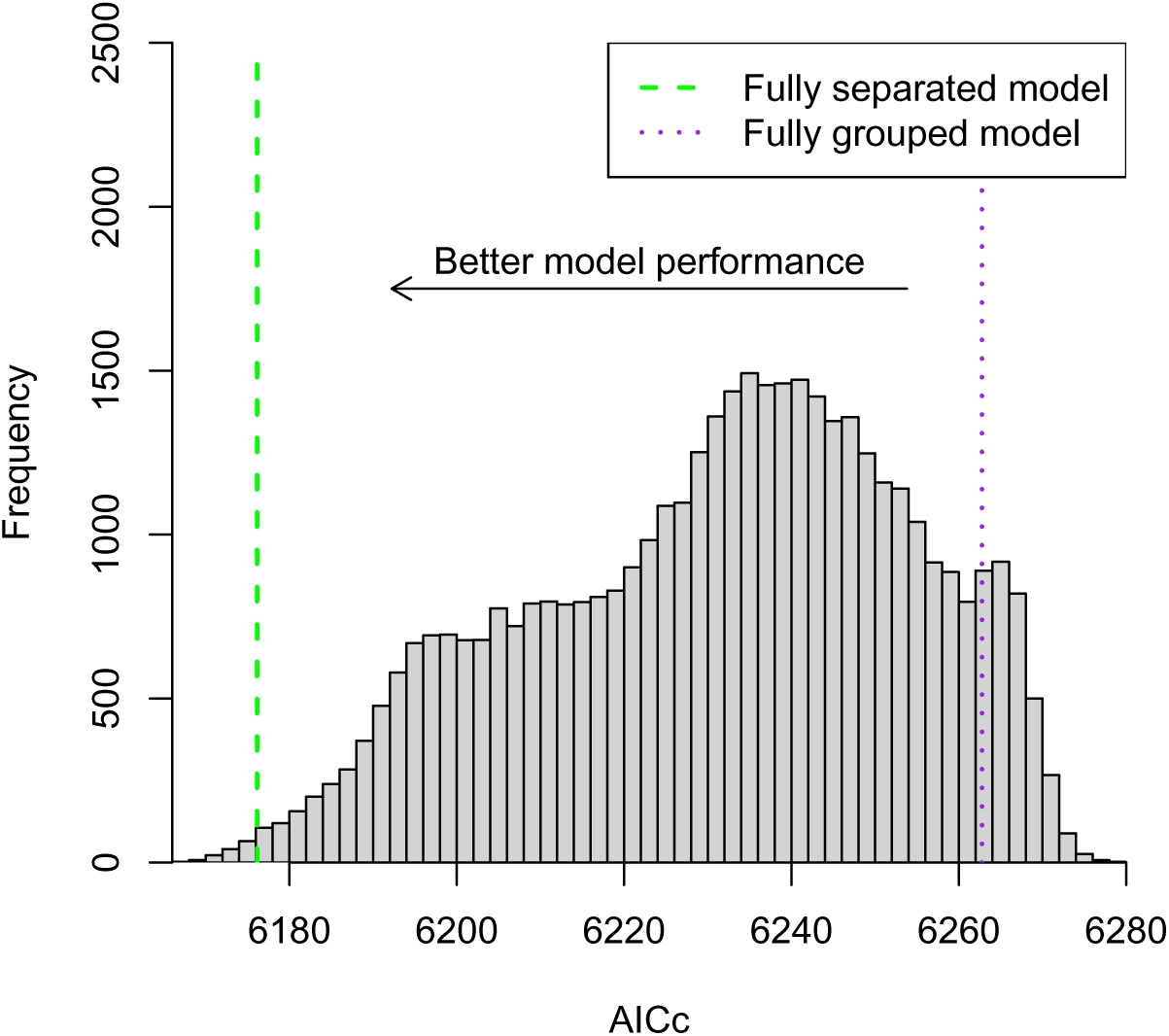
Histogram of the AICc values of Beverton–Holt model fits with all possible groupings of interaction coefficients. The dashed green line shows the AICc value (6176.14) of the ungrouped model in which all competition coefficients are fitted independently. The dotted purple line shows the AICc value (6262.76) of the fully grouped model in which every pair of species shares a single interaction coefficient.

The “best” model, as indicated by AICc, was not the best with any surety; its Akaike weight indicated a probability of only 0.096 that it was the best. Figure 2 shows the groupings and interaction coefficients of this “best” model. The ungrouped model had an AICc just 9.35 units from the best-performing model, but the total Akaike weight assigned to models performing better than the ungrouped model was 0.893; this is the probability that some grouped model performed better than the ungrouped model.

**Figure 2:**
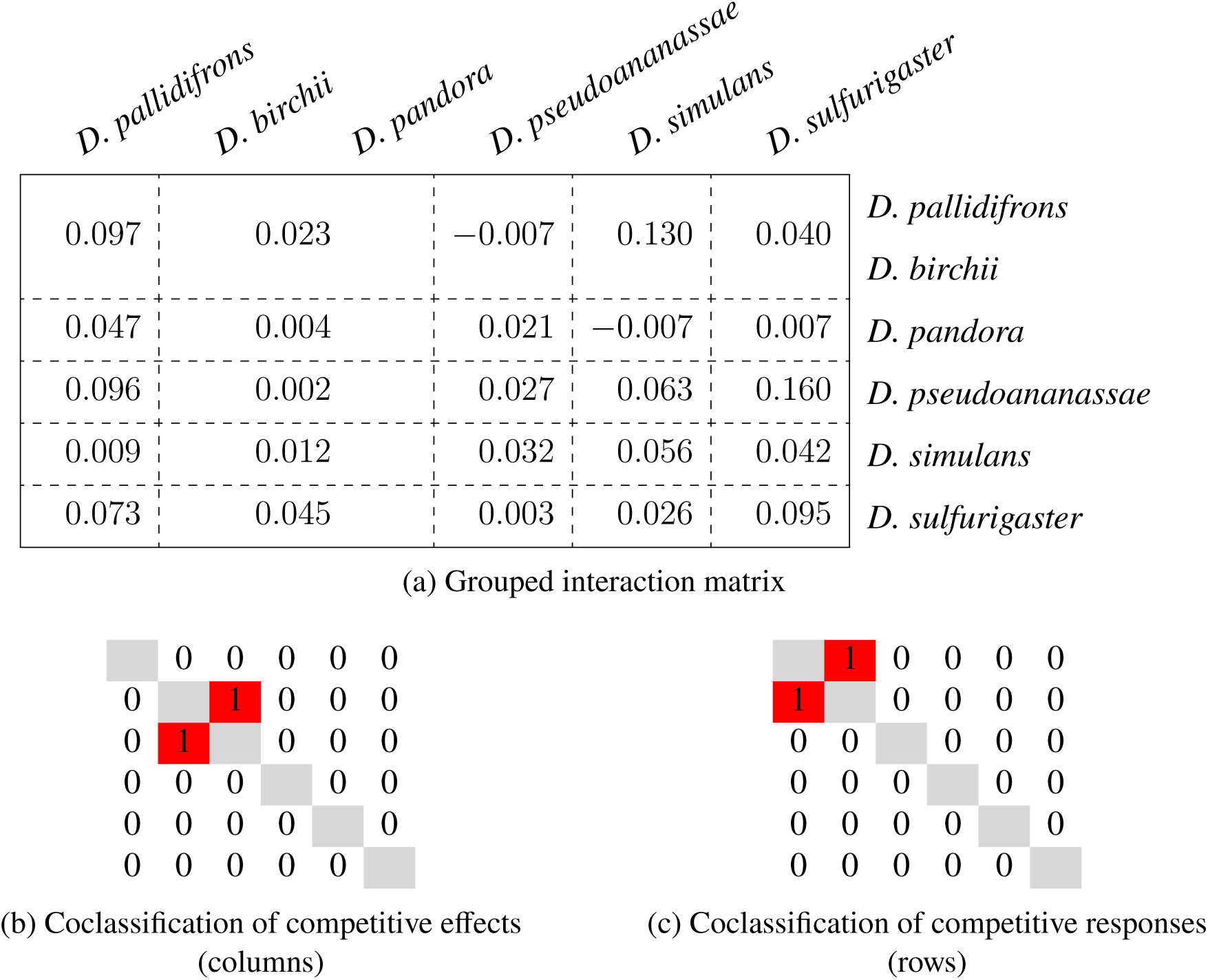
The groupings and competition coefficients of a grouped Beverton–Holt model fitted to the six studied *Drosophila* species, where results are for the single model with the best AICc (6166.790). (a) We show the inferred matrix of competition coefficients, where a competition coefficient in a block represents the same coefficient being used for multiple interactions (per the identified groupings). (b) and (c) We show the coclassification matrices derived from the groupings of the columns and rows, respectively. Species have been reordered so that species that were grouped together appear adjacent, and the ordering is the same in all of (a), (b), and (c).

Two other models lay within 2 AICc units of the “best” model, and thus, by the traditional rule of thumb, would also be considered viable candidates as the “best” model (Cavanaugh and Neath 2019). The three “best” models had Akaike weights of 0.096, 0.068, and 0.039, for a total of 0.203. These models had, respectively, a number of row and column groups equal to 5 by 5, 4 by 4, and 4 by 5. Thus, they had respectively 32, 23, and 27 parameters.

Considering only models in which the rows and columns were grouped using the same grouping, the AICc values ranged from 6172.26 to 6264.85; neither the best nor worst overall models had equivalently grouped rows and columns. The total Akaike weight assigned to these models was 0.007; this was more than expected by chance (0.005), but not significantly so (Monte Carlo permutation test, *p* = 0.183). This indicates that the best-performing model was not especially likely to be one that grouped the rows and columns equivalently.

### 3.2 Coclassification of species

The highest probability of coclassification in either the row or column groupings was 64.4 % (fig. 3). While this means that no pair of species could be confidently be said to be coclassified, the results from the simulated data (fig. S4) show that this is no guarantee that the species did not belong to the same group either. In all, some pairs of species were far more likely to be coclassified than others, and many pairs of species were quite certain not to be coclassified, suggesting that there exists some structure underpinning the best groupings.

**Figure 3:**
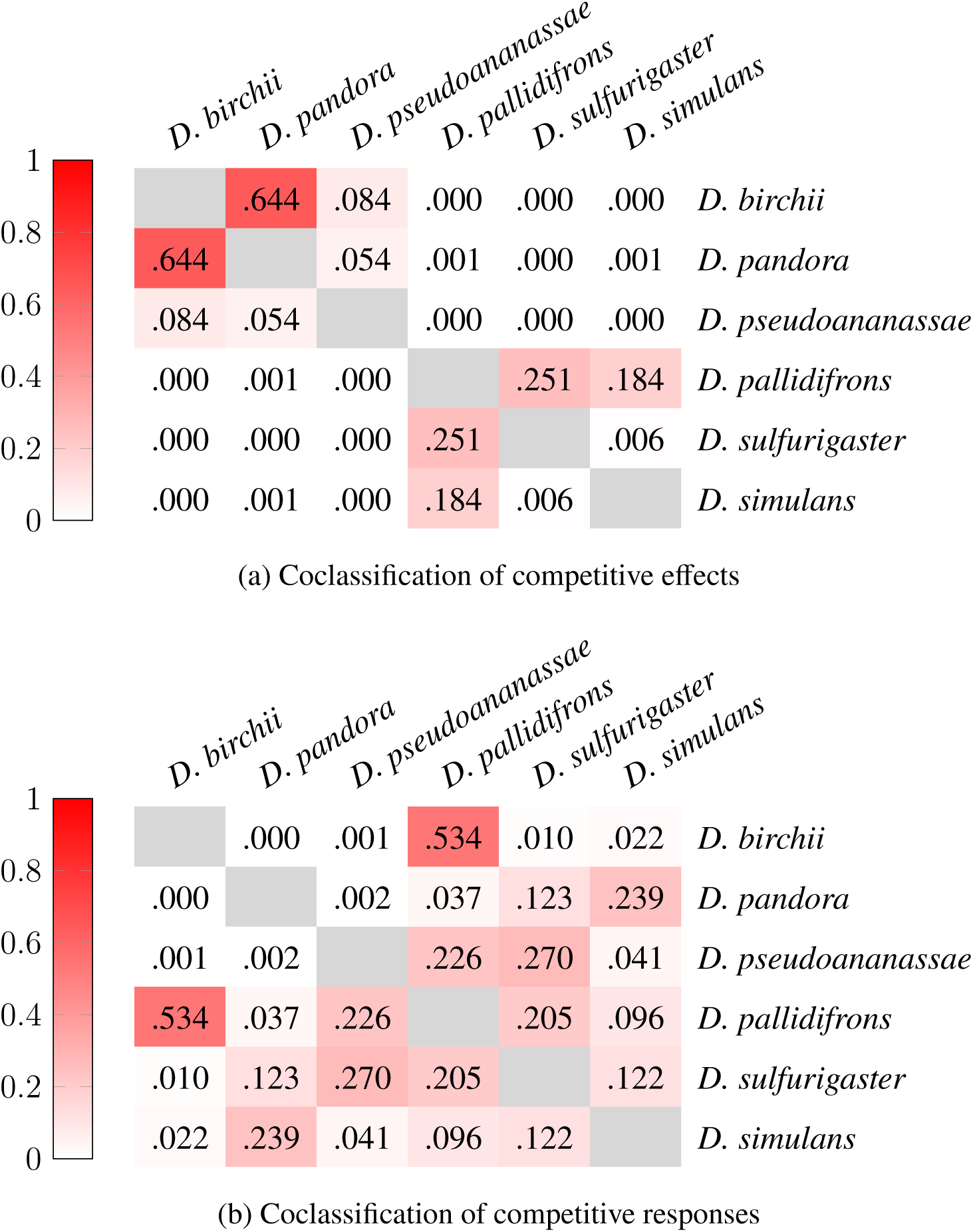
Coclassification matrices of the Beverton–Holt competitive effects and responses of the six studied *Drosophila* species, weighted by Akaike weights. The cell for a pair of species represents the probability that the best grouping, as determined by the Akaike weights of the grouped models, places those two species in the same group. (a) We show the grouping of species’ competitive effects upon other species (columns of the interaction matrix). (b) We show the grouping of species’ responses to competition (rows of the interaction matrix).

The structure of the coclassification matrices was very different in the row groups than in the column groups. That is, most pairs of species which had a high probability of being coclassified in one grouping had a very low probability of being coclassified in the other grouping. In particular, in the column groupings (groupings of competitive effects), *D. birchii* was frequently coclassified with *D. pandora* but was almost never coclassified with *D. pallidifrons*, while in the row groupings (groupings of competitive responses) this was completely reversed. Overall, more coclassification occurred in the row (response) groupings than in the column (effect) groupings. A Mantel test of the coclassification matrix for the rows against that for the columns showed that there is no significant correlation between the two (*r*_M_ = −0.243, *p* = 0.827). PERMANOVA showed that neither the row nor column coclassification matrix was significantly explained by species body size (column: *F*_1,4_ = 1.973, *p* = 0.074; row: *F*_1,4_ = 0.589, *p* = 0.860). Testing the correlation of average competitive effects and competitive responses (column- and row-averaged interaction coefficients from the ungrouped model) with species body size, competitive effects are positively correlated with body size (*r* = 0.767, *p* = 0.038), but competitive responses are not (*r* = −0.056, *p* = 0.542).

## 4 Discussion

We have shown that it is possible to obtain a better performing model of species interactions by grouping the interaction coefficients of species *a posteriori*, and hence reducing the number of free parameters in the model. Indeed, the results show that at least one grouping of species was likely to give a better performing model (probability 0.893) than leaving them ungrouped. This result contrasts with the results of Buche et al. (2025), who found that the degree of grouping (species, family, or functional group) did not significantly affect their model’s performance. This may be because, while they tested for the appropriate *level* of grouping, their assignment of species to groups was always decided *a priori*. This suggests that using a data-informed approach to classifying species into groups is important to gaining maximal benefit from grouping.

While grouping did improve model performance, which exact grouping would give the best-performing model for the data set studied here could not be determined with confidence. This result stands in contrast to the observation that species sometimes form clear groups, be they the functional groups so often used (Blondel 2003; Kirwan et al. 2009; Boulangeat et al. 2012), or taxonomic groups per a relationship between phylogeny and ecological niche (Godoy et al. 2014; Wilcox et al. 2018). It also contrasts with previous studies that have succeeded in finding clear groups in other ecological contexts, such as in food webs (Krause et al. 2003; Allesina and Pascual 2009) or plant–pollinator networks (Olesen et al. 2007).

One possible explanation for our results derives from our method optimising for predictive power, rather than for maximum model simplification (i.e. maximum grouping). Using BIC, rather than AIC, would likely have grouped species further since BIC—in searching for the “true” model rather than the model with the greatest predictive power—prefers to simplify models further than AIC (Wagenmakers and Farrell 2004; Aho et al. 2014). Another possible explanation for our results is that species simply do not group to the same extent in a quantitative competitive context as in binary interaction networks. That said, the results suggest that this explanation is unlikely as grouping was useful for improving model performance. In the present case, we cannot rule out the possibility that a lack of clarity in the groupings is instead a product of the study system. All the species in this study were members of the same genus, *Drosophila*, and as such were all rather similar. A study system with species spanning multiple functional groups, or a broader taxonomy, might have shown less equivocal groupings.

Alas, the brute force method we employed in this study has a very poor algorithmic complexity, complicating any application of this approach to studies with more than seven or eight species (there are (*B*_7_)^2^ = 769 129 possible models with 7 species and (*B*_8_)^2^ = 17 139 600 models with 8 species). Nonetheless, our results might provide motivation to develop an improved method with which to test for grouping of species interactions among a larger set of species. For example, reversible-jump Markov chain Monte Carlo methods would allow selection among many possible groupings without requiring an exhaustive search of every such grouped model (Green 1995). Alternatively, a group lasso (Yuan and Lin 2006) might be able to select the best among the possible groupings, while penalising models with too many parameters in the same fashion as AICc. Further, a sparse-group lasso (Simon et al. 2013) may be able to combine this the lasso approach with a similar sparse matrix approach to that employed by Weiss-Lehman et al. (2022) and Buche et al. (2025). Instead, it may simplify matters to adopt a mixed-membership approach in which a species can belong to more than one group (Airoldi et al. 2008), reminiscent of the community-level drivers of Ovaskainen et al. (2017).

Coclassification between most pairs of species was quite low. This suggests that several species would best stand on their own, ungrouped with other species. This is contrary to the results of Weiss-Lehman et al. (2022) and Buche et al. (2025), who concluded through sparse matrix modelling that only a few species had competitive effects differing from the mean. We would have expected a similar outcome in our results to manifest as most species belonging to the same group (for the grouping of the competitive effects, at least), with only a few species falling outside of that group. It seems worth noting, however, that even a relatively low degree of grouping can result in a substantial reduction in the number of parameters in the model. For example, although the model with the lowest AICc only grouped one pair of species in the rows and one pair in the columns, it reduced the number of parameters in the model to 32, compared to 43 parameters in the ungrouped model. Another candidate “best” model (ΔAIC *<* 2) had as a few as 23 parameters. When viewed in the context of the experimental data studied here, this implies that one could hypothetically have conducted a response–surface experimental design with scarcely half as many treatments and still achieved equivalent predictive performance. Furthermore, the quadratic scaling of the number of interaction coefficients means that the added benefit of even small amounts of grouping would only continue to grow as the total number of species in the system increased.

While our results could not definitely identify a single best grouping, one thing that was clear was that the model-supported groupings differed for the rows and columns. That is, species that are similar in their response to competition may not be similar in the competitive effects they have on other species, and vice versa. In hindsight, this should not be overly surprising since it is known that different traits govern species competitive effects than govern their responses (Goldberg 1990; Goldberg and Landa 1991); for example shading effect and shade tolerance in plants. In this system, average competitive effect was correlated with body size, suggesting that body size is a trait governing competitive effect, though it was unlikely to be the only trait influencing competitive effect as it did not suffice to explain the grouping in competitive effect. However, body size had no correlation with competitive response, nor did it explain the grouping in competitive response, suggesting that a different trait is responsible for determining response (the original authors did not record any other species traits that could be tested). Yet the assumption that species should group equivalently in regards to competitive effects and responses is frequently made, often implicitly and without comment (e.g. Ives et al. 2003; Martorell and Freckleton 2014; Gross and Edmunds 2015). Our results recommend more explicit questioning, and even testing, of this assumption when building future species-interaction models.

We observed some intransitivity in the two coclassification matrices. For example, in the column groupings, *D. pallidifrons* had a relatively high probability of being coclassified with *D. birchii*, and a substantial probability of being coclassified *D. pseudoananassae*, yet *D. birchii* and *D. pseudoananassae* had almost no probability of being coclassified themselves. This suggests that there may not be a single “best” grouping of the species, but rather two or more quite distinct and mutually exclusive groupings of similar validity. A single trait like body size might therefore not be able to capture the multivariate nature of species’ competitive niches (Blonder 2018). This could indicate that a Bayesian approach is worth exploring in the future, in which the outcome is not a definitive grouping but a distribution of valid groupings. Alternatively, this may suggest further value in the mixed-membership approach mentioned above.

## 5 Data availability

All raw observation data used in the manuscript are archived on Zenodo https://doi.org/10.5281/zenodo.4736655 and GitHub https://github.com/jcdterry/TCL_DrosMCT/tree/1.0.0. All code used to perform the analyses is also archived on Zenodo https://doi.org/10.5281/zenodo.17493256 and GitHub https://github.com/pyrrhicPachyderm/interaction-partitioning/tree/ms1-v4.

## Acknowledgements

We would like to thank Hao Ran Lai for feedback on a draft of the manuscript. We would also like to thank Chris Terry for a number of helpful comments during peer review. DBS is also grateful for the support of the Marsden Fund Council, from New Zealand Government funding (grant 16-UOC-008).

## Authors’ contributions

DBS conceptualised the investigation. CRPB developed and implemented the analysis, with input and feedback from DBS. CRPB wrote the first manuscript draft. Both authors contributed to manuscript edits.

