## Supplementary material for "Simplifying species-interaction models by grouping parameters: optimal groupings differ between effects and responses"

### S1 Supplementary methods

To assess the ability of our method to discern groups, we applied the method to simulated data with known group structure. We used four species in the simulated data, divided into two groups of two species each: the minimum number necessary to have multiple groups each containing multiple species. In each set of simulated data, the alpha values for species belonging to the same group were identical, but we manually adjusted (i) the difference between the average alpha values of the two row groups and (ii) an independent difference between the average alpha values of the two column groups, thus forming four blocks of alpha values in the matrix. The differences between the two groups were set as a proportion, from $0\,\%$ to $150\,\%$, of the overall average alpha value. That is, letting the overall average alpha value be $\bar{\alpha}$, and the proportional row and column differences be $\Delta_r, \Delta_c \in [0, 1.5]$, the alpha values of the four groups were:

$$\alpha_{1,1} = \alpha_{1,2} = \alpha_{2,1} = \alpha_{2,2} = \bar{\alpha}\left(1 - \frac{\Delta_r}{2} - \frac{\Delta_c}{2}\right),$$
$$\alpha_{1,3} = \alpha_{1,4} = \alpha_{2,3} = \alpha_{2,4} = \bar{\alpha}\left(1 - \frac{\Delta_r}{2} + \frac{\Delta_c}{2}\right),$$
$$\alpha_{3,1} = \alpha_{3,2} = \alpha_{4,1} = \alpha_{4,2} = \bar{\alpha}\left(1 + \frac{\Delta_r}{2} - \frac{\Delta_c}{2}\right),$$
$$\alpha_{3,3} = \alpha_{3,4} = \alpha_{4,3} = \alpha_{4,4} = \bar{\alpha}\left(1 + \frac{\Delta_r}{2} + \frac{\Delta_c}{2}\right).$$

All other components of the simulated data were designed to emulate those of the real dataset used in the main text. The growth rate (which for simplicity was fixed to be equivalent for all species in the simulated data), the overall average alpha value, and the dispersion parameter were taken as the average values of those parameters in the ungrouped model fitted to the real data. The design matrix of the experiments was also based on that of the real data: each single species in densities of 6, 18, and 30 individuals, and each pair of species in densities of (6,6), (6,12), (6,18), (6,24), (12,6), (18,6), or (24,6) individuals, with each replicated three times. The mean numbers of offspring were then calculated using the same Beverton–Holt model as was fit to the real data, and this mean and the aforementioned dispersion parameter were used to draw negative-binomially distributed random observations.

We simulated $360$ sets of data: with the differences between the row and column groups each independently set to between $0\,\%$ and $150\,\%$ of the overall average alpha value, in intervals of $25\,\%$, and with ten replicate datasets for each case. To each dataset, we applied the same model-fitting procedure as applied to the real data: fitting a model for each possible combination of row and column groupings, calculating the AICc and corresponding Akaike weight for each such model, and using the Akaike weights to generate row and column coclassification matrices for the species in each simulated dataset. The entries from the resulting coclassification matrices were divided into within-group and between-group entries, based on the groups known to exist in the simulated data. A plot of the resulting coclassification values, as a function of the known difference between group alpha values, is shown in fig. S4, and the results are discussed in the main text.

### S2 Formalisms

The following establishes some of the concepts used in main text more formally.

#### S2.1 Groupings

In mathematics, the groupings we refer to in the main text are formally known as partitions of a set. A partition of a set is the division of its elements into a set of non-empty subsets, so that each element of the original set appears in exactly one subset. If we use the integers $\{1, 2, 3, 4, 5, 6\}$ to represent the six species, a few example partitions are $\{\{1\}, \{2\}, \{3\}, \{4\}, \{5\}, \{6\}\}$ (when each species belongs to its own group, yielding $6$ groups), $\{\{1, 2, 3, 4, 5, 6\}\}$ (when all species belong to the same group, yielding $1$ group), and $\{\{1, 3\}, \{2, 4, 6\}, \{5\}\}$ (an intermediate case with $3$ groups). The cardinality $|P|$ of a partition $P$ is the number of groups in the partition.

In the following, we will use $P_{\mathrm{g}}$ to refer to the partition with all species in one group (the fully grouped case), and $P_{\mathrm{s}}$ to refer to the partition with all species in different groups (the fully separated case):

$$P_{\mathrm{g}} = \{\{1, \ldots, n\}\}, \tag{S1}$$

$$P_{\mathrm{s}} = \{\{1\}, \ldots, \{n\}\}. \tag{S2}$$

We also define:

$$t_{ij}(P) = \begin{cases} 1, & \text{if } i \text{ and } j \text{ are in the same subset within } P \\ 0, & \text{otherwise.} \end{cases} \tag{S3}$$

We will use $\mathcal{P}$ to refer to the set of all possible partitions of $n$ species.

#### S2.2 Models

Each model we consider is defined by two partitions: one for the rows of the interaction matrix, and one for the columns. Thus, we can denote a model as $m_{P_a P_b}$, where $P_a$ is the partition of the species in the rows, and $P_b$ the partition of the species in the columns. Formally, for a model $m_{P_a P_b}$, $\alpha_{i_1 j_1} \equiv \alpha_{i_2 j_2}$ if $t_{i_1 i_2}(P_a) = 1$ and $t_{j_1 j_2}(P_b) = 1$. Model $m_{P_a P_b}$ thus has $k(m_{P_a P_b}) = n + |P_a| \cdot |P_a| + 1$ parameters: $n$ growth rates, $|P_a| \cdot |P_a|$ interaction coefficients, and $1$ overdispersion parameter.

#### S2.3 AIC & Akaike weight

The AIC value for a model depends upon the maximised value of the likelihood function of the model, and the number of parameters in the model:

$$\text{AIC}(m_{P_a P_b}) = 2k(m_{P_a P_b}) - 2\log\left(\hat{L}(m_{P_a P_b})\right), \tag{S4}$$

where $k$ is the number of parameters in the model (defined above), and $\hat{L}$ is the maximised value of the likelihood function for the model, given the data. AICc contains a correction term for small sample sizes:

$$\text{AICc}\left(m_{P_aP_b}\right) = \text{AIC}\left(m_{P_aP_b}\right) + \frac{2k\left(m_{P_aP_b}\right)^2 + 2k\left(m_{P_aP_b}\right)}{l - k\left(m_{P_aP_b}\right) - 1}, \tag{S5}$$

where $l$ is the sample size of the data. To convert AICc to Akaike weights, we must first calculate $\Delta$AICc for each model:

$$\Delta\text{AICc}\left(m_{P_aP_b}\right) = \text{AICc}\left(m_{P_aP_b}\right) - \min_{P_u\in\mathcal{P}, P_v\in\mathcal{P}} \text{AICc}\left(m_{P_uP_v}\right). \tag{S6}$$

Then the Akaike weight $w$ for each model is:

$$w\left(m_{P_aP_b}\right) = \frac{\exp\left(-\frac{1}{2}\Delta\text{AICc}\left(m_{P_aP_b}\right)\right)}{\sum_{P_u\in\mathcal{P}, P_v\in\mathcal{P}} \exp\left(-\frac{1}{2}\Delta\text{AICc}\left(m_{P_uP_v}\right)\right)}. \tag{S7}$$

Fitting their interpretation as probabilities, the sum of the Akaike weights for all the models considered is $1$.

#### S2.4 Coclassification matrices

With the Akaike weights in hand, the values of the coclassification matrices can be defined. The set of models yields two coclassification matrices: one for the row groupings, which we shall denote $C_{\text{row}}$, and one for the column groupings, $C_{\text{col}}$. The value of each cell of the row coclassification matrix is:

$$c_{\text{row},ij} = \sum_{P_a\in\mathcal{P}, P_b\in\mathcal{P}} t_{ij}\left(P_a\right) w\left(m_{P_aP_b}\right), \tag{S8}$$

and the value of each cell of the column coclassification matrix is similarly:

$$c_{\text{col},ij} = \sum_{P_a\in\mathcal{P}, P_b\in\mathcal{P}} t_{ij}\left(P_b\right) w\left(m_{P_aP_b}\right). \tag{S9}$$

Since a species tautologically appears in the same subset of a partition as itself, the diagonal elements of the coclassification matrices are $1$, i.e. $c_{\text{row},ii} = c_{\text{col},ii} = 1\ \forall i$.

##### S2.4.1 Reference values

The main text refers to three useful reference values for the off-diagonal elements coclassification matrices. We show the formal derivation of these here.

First, the neutral model approach, where all species are considered to be equivalent, and thus in the same group as one another. Here:

$$w\left(m_{P_aP_b}\right) = \delta_{P_aP_\text{g}}\delta_{P_bP_\text{g}}, \tag{S10}$$

where $\delta_{ij}$ is the Kronecker delta, i.e. $1$ if $i = j$ and $0$ otherwise. Thus, in the neutral model case, $c_{\text{row},ij} = c_{\text{col},ij} = 1\ \forall i, j$.

Second, if every species is expected to be unique, and thus in its own group, then:

$$w\left(m_{P_aP_b}\right) = \delta_{P_aP_\text{s}}\delta_{P_bP_\text{s}}. \tag{S11}$$

Thus, in this case, $c_{\text{row},ij} = c_{\text{col},ij} = 0\ \forall i \neq j$.

Finally, in the case where each possible grouping is considered to be equally likely, noting that there are $B_n$ possible partitions of $n$ species:

$$w\left(m_{P_aP_b}\right) = \frac{1}{B_n^2}. \tag{S12}$$

Thus, in this case, $c_{\text{row},ij} = c_{\text{col},ij} = \frac{B_{n-1}}{B_n}\ \forall i \neq j$. To see why the numerator is $B_{n-1}$, note that $\sum_{P_a \in \mathcal{P}} t_{ij}\left(P_a\right) = B_{n-1}$, i.e. there are $B_{n-1}$ partitions of $n$ elements that place two given elements in the same group. This is because, with two elements necessarily placed in the same group, they may be treated as a single element of a set with one fewer elements, which may thus be partitioned in $B_{n-1}$ possible ways.

#### S2.5 Other important statistics

The main text also references two other important statistics that can be calculated from the set of Akaike weights for each model. We provide their formal definitions here.

The first is the total Akaike weight of models with a lower AICc than the entirely ungrouped model:

$$\sum_{P_a \in \mathcal{P}, P_b \in \mathcal{P}} w\left(m_{P_a P_b}\right)\left[w\left(m_{P_a P_b}\right) > w\left(m_{P_\mathrm{s} P_\mathrm{s}}\right)\right], \tag{S13}$$

where $[S]$ is the Iverson bracket of $S$, i.e. $1$ if the statement $S$ is true, and $0$ otherwise. As the Akaike weight of a model is the probability that it is the best-performing model among those assessed, this sum gives the total probability that there is some model which performs better than the entirely ungrouped model.

The second important statistic is the total Akaike weight of models that used the exact same grouping for the rows as for the columns:

$$\sum_{P_a \in \mathcal{P}} w\left(m_{P_a P_a}\right). \tag{S14}$$

This gives the probability that the best-performing model was one which grouped species equivalently in their competitive effects and competitive responses.

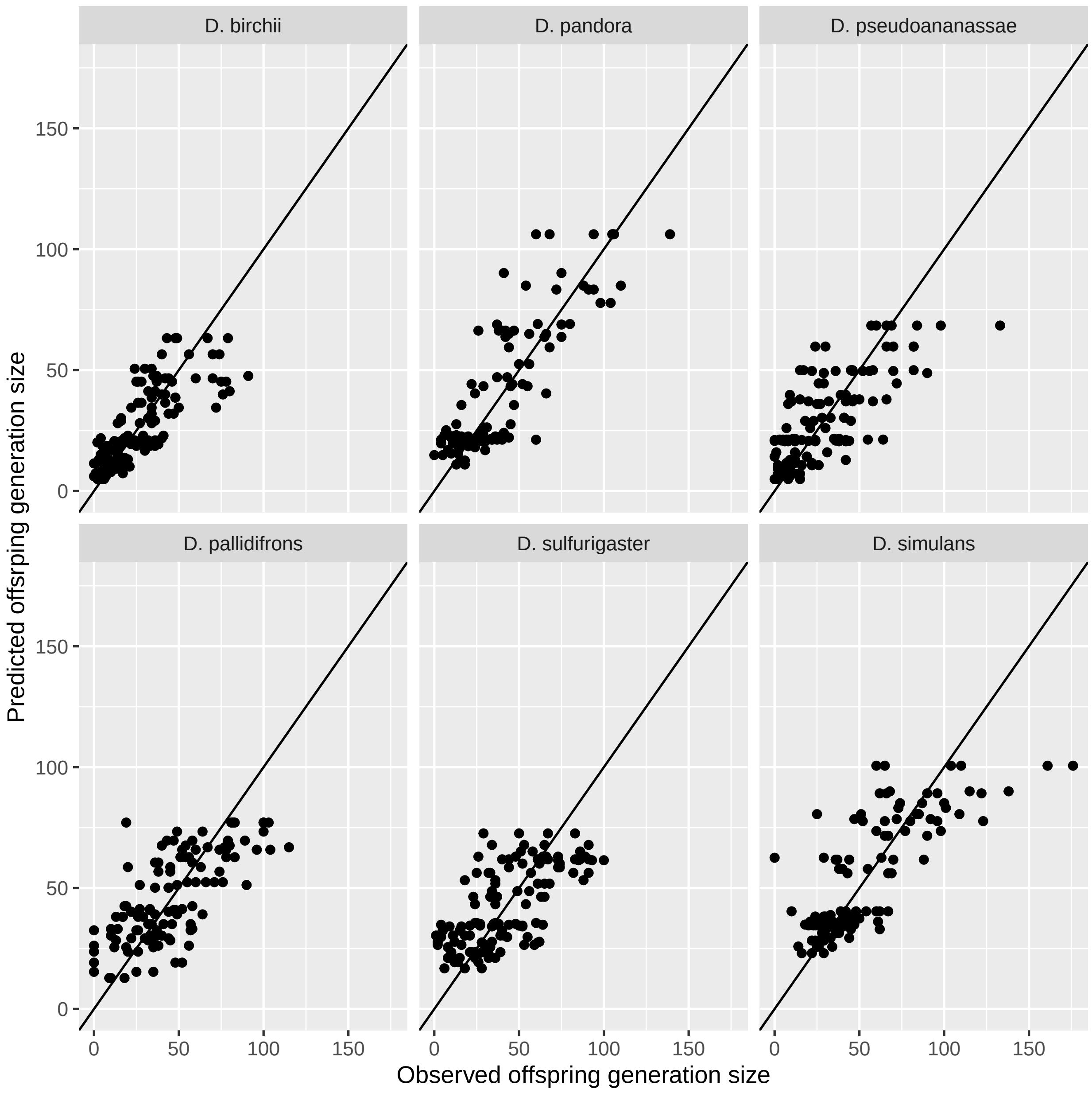


Figure S1: Plot of model fit using the best performing model (as indicated by AICc). The black line shows $y = x$.

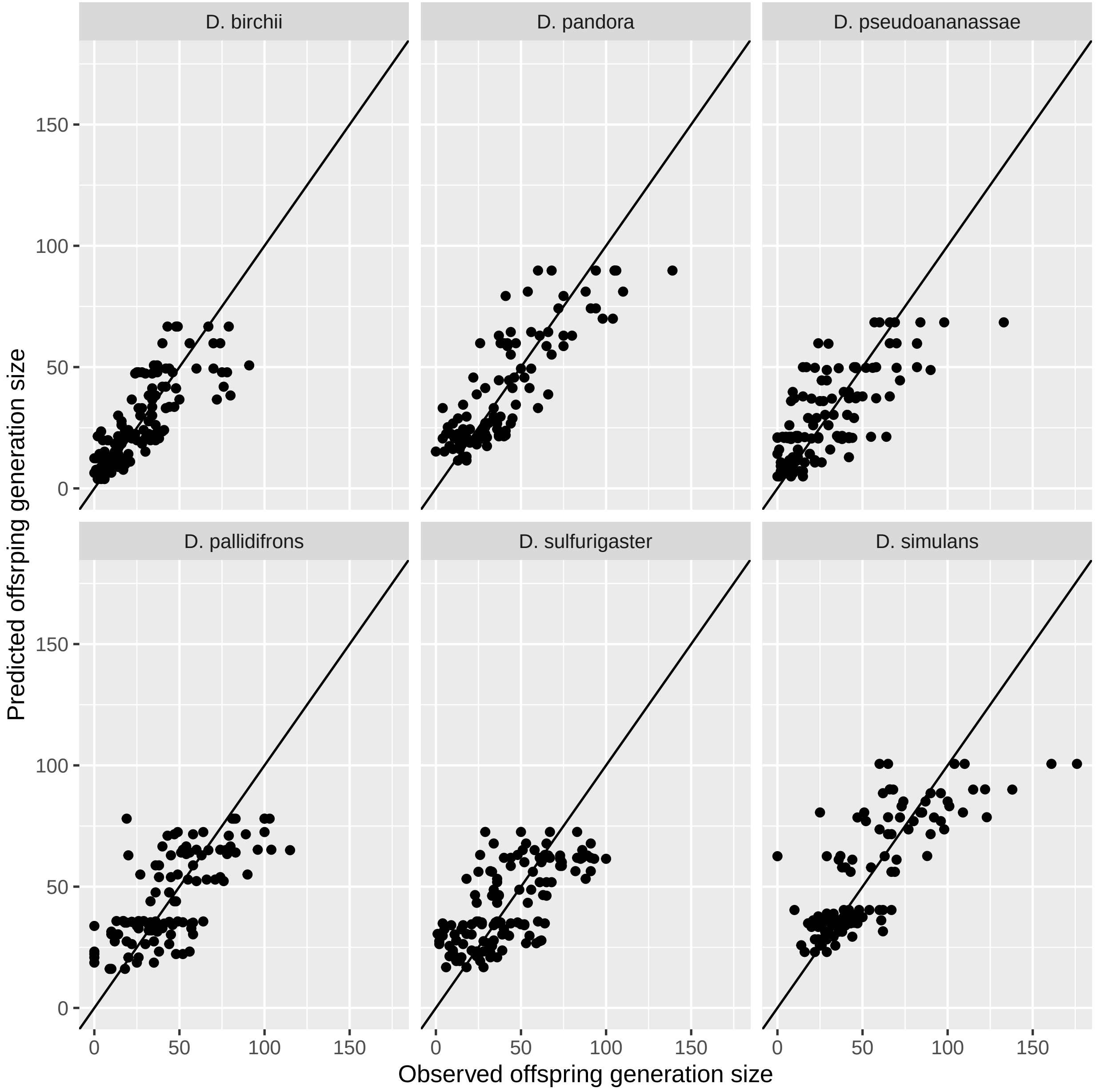


Figure S2: Plot of model fit using the fully separated model. The black line shows $y = x$.

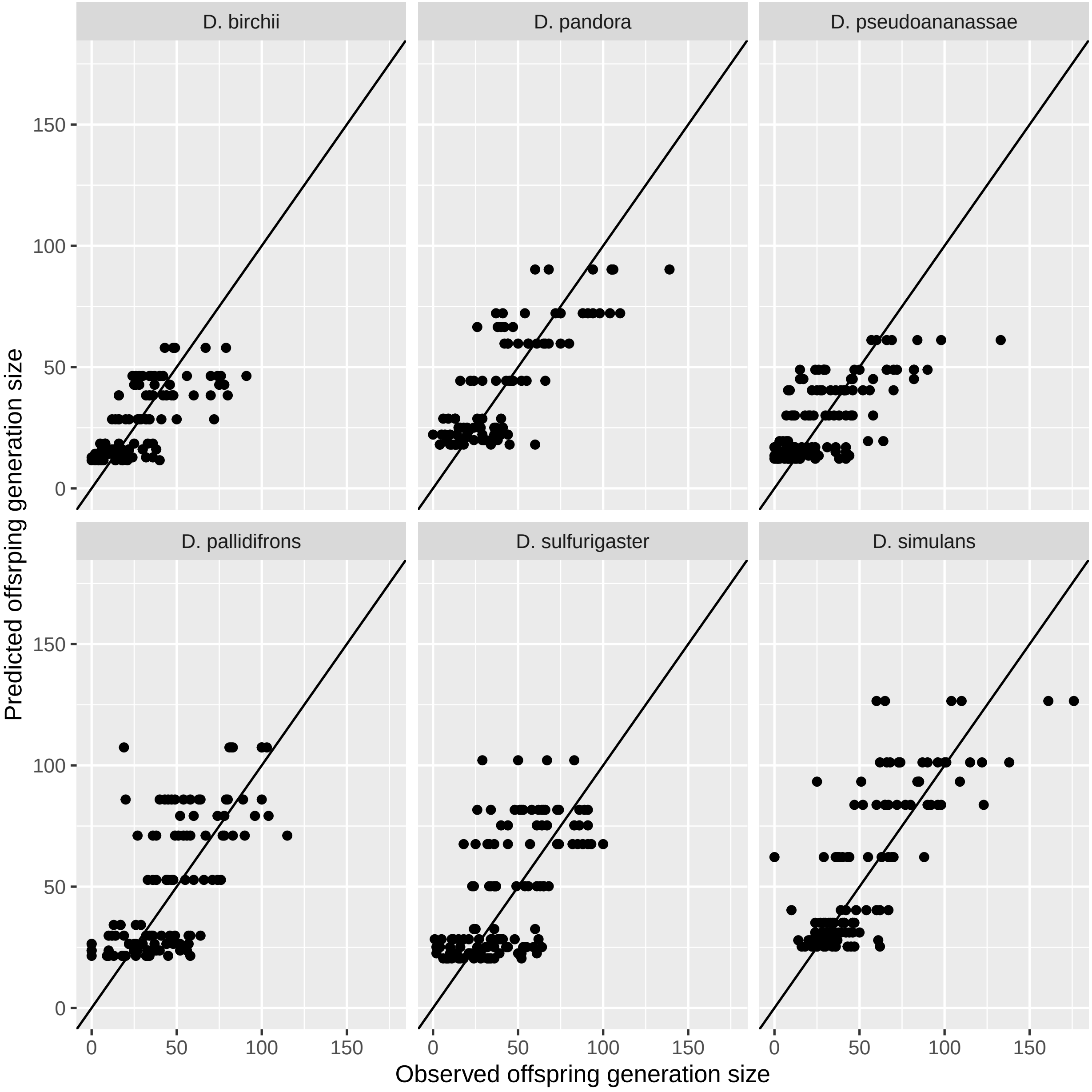


Figure S3: Plot of model fit using the fully grouped model. The black line shows $y = x$.

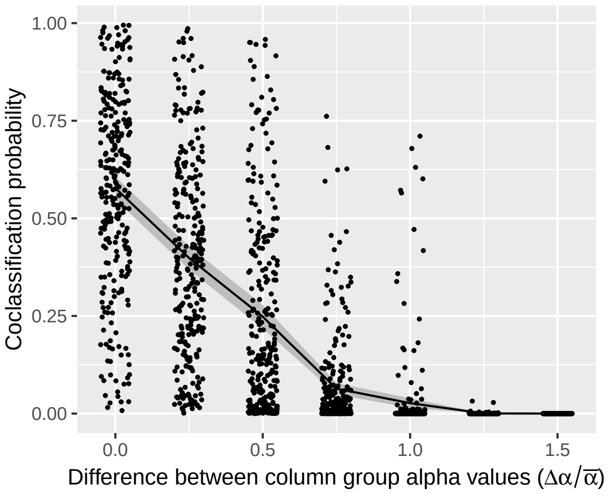


(a) Competitive effect (column) coclassification between groups

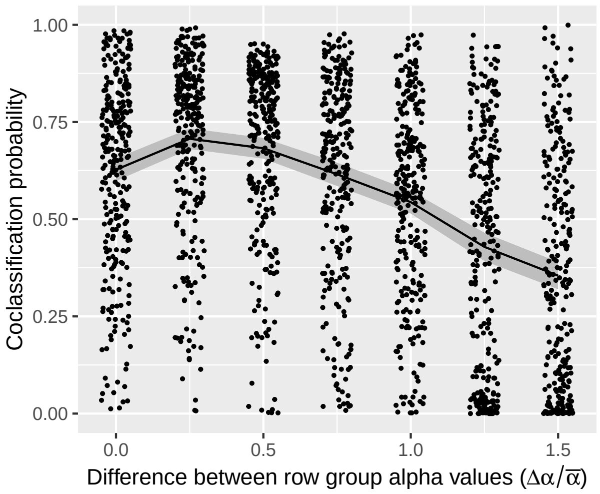


(b) Competitive response (row) coclassification between groups

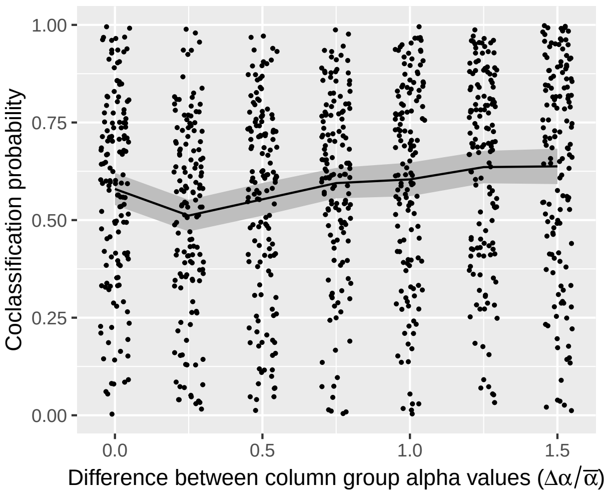


(c) Competitive effect (column) coclassification within groups

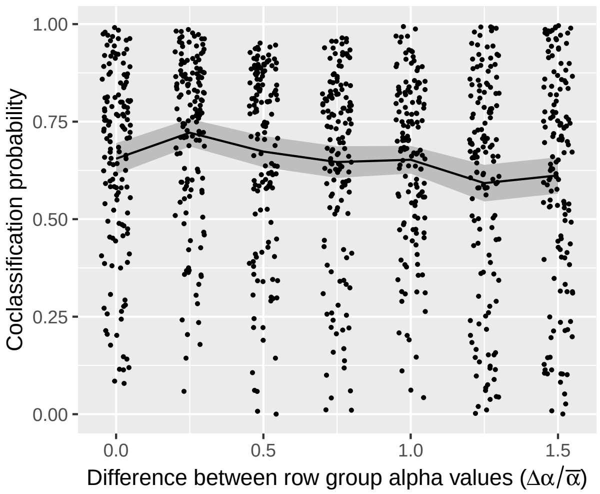


(d) Competitive response (row) coclassification within groups

Figure S4: Plot of coclassification results from simulated data with known groups. Each point represents a single cell from a coclassification matrix generated from a dataset. Points have been horizontally jittered for visibility. The line shows mean coclassification values, while the shaded area shows the $95\,\%$ confidence interval for the mean. (a) and (b) show coclassification probabilities between pairs of species known to be in different groups in the generated data, while (c) and (d) show coclassification between pairs of species known to be in the same group. (a) and (c) show coclassification in the competitive effect (column) groups, against the known difference between the alpha values of the column groups, while (b) and (d) show the same for the competitive response (row) groups, against the known difference between the row groups.
